# Comparable neural and behavioural performance in dominant and non-dominant hands during grasping tasks

**DOI:** 10.1101/2024.12.09.627653

**Authors:** Balasubramanian Eswari, Sivakumar Balasubramanian, SKM Varadhan

## Abstract

Hand dominance has long been associated with differences in neural control and motor performance, with the dominant hand typically exhibiting better coordination in reaching tasks. However, the extent to which this dominance influences performance in finger force control remains unclear. This study aimed to examine the behavioural and neural features of the dominant and non-dominant hands during grasping and lifting tasks in healthy young adults, focusing on the synergy index, EEG band power, and EEG–EMG coherence as key measures. Twenty right-handed adults (mean age = 26.95, STD = 2.68) participated in this study. Participants engaged in an experimental task where they grasped a handle for the initial five seconds, followed by lifting and holding it for an additional five seconds. There were two task conditions: fixed (thumb platform secured) and free (thumb platform movable). They performed 25 trials with both the dominant and non-dominant hands in the two task conditions, with the order of trials and hands block randomized to eliminate potential order effects. Contrary to the dynamic dominance hypothesis, we found statistical equivalence in the synergy index, EEG band power, and EEG–EMG coherence between the dominant and non-dominant hands across both fixed and free task conditions. These findings suggest that both hands are capable of achieving similar levels of performance in tasks emphasizing steady-state force maintenance, despite the typical advantages of the dominant hand in other motor tasks. While task-dependent modulations in behavioural and neural features were observed due to changes in friction, these adjustments were non-different between the dominant and non-dominant hands.

## Introduction

Human hands are highly dexterous, capable of manipulating objects with both the dominant and non-dominant hands. Previous studies on reaching tasks have revealed notable differences between the dominant and non-dominant limbs in handling movement trajectories and accomplishing precise endpoint locations [1], [2]. According to the dynamic dominance hypothesis, the dominant (DOM) hand, typically controlled by the left hemisphere in right-handed individuals, demonstrates precision and superior trajectory control. This hand is used for writing, grasping a pen, and performing tasks that necessitate fine motor skills [3]. In contrast, the non-dominant (NDOM) hand, governed by the right hemisphere, primarily serves as a stabilizer. It ensures that objects remain secure, applies appropriate force, and maintains steady, controlled movements [4]. These functional differences between the dominant and non-dominant hands highlight the complicated balance of control exerted by each hemisphere during motor tasks. Considerable research supports the general notion that the contralateral hemisphere of the cortex predominantly controls hand movements, emphasizing the brain’s critical role in coordinating motor functions [5], [6], [7].

Although the brain’s two hemispheres have similar anatomical structures, they function asymmetrically: the left hemisphere primarily manages language and analytical tasks, while the right hemisphere is more involved in spatial and creative functions [8]. This functional asymmetry is particularly significant in motor control. The dominant hemisphere’s primary motor cortex (M1) has a larger hand area with more connected movement representations, allowing for better learning and coordination of motor skills [9], [10]. This enhanced connectivity supports superior motor practice and dexterity in the dominant hand compared to the non-dominant hand. Task-induced muscle evoked potential during the pegboard task showed increased corticomotor excitability for gripping muscles in both hands, but only the dominant hemisphere showed reduced inhibition and links to motor learning. This suggests hemispheric differences in controlling skilled hand movements [11].

The meticulous regulation of force depends on the sophisticated coordination between the central nervous system and hand musculature, emphasizing the importance of neuromuscular communication. Corticomuscular coherence (CMC) in the 15–30 Hz frequency range indicates the functional connection between the cortex and muscles through EEG and EMG signals [12], [13]. In healthy individuals, CMC magnitude increases as EMG signals become strong [14] and also during maximum voluntary contractions (MVC) [15]. Beta-range corticomuscular coherence showed a significant difference between the left and right hands during gripping tasks [16]. Previous studies focused on isometric force production or grip force control in finger-pressing tasks have shown varying CMC magnitude between the hands in some tasks, while demonstrating no differences in some other tasks [16], [17], [18].

In motor control, multi-digit grasping requires the coordination of grip and tangential forces to successfully complete tasks, relying on precise control and adjustment of finger forces. In this context, synergy refers to the coordinated adjustment of finger forces and moments produced by each finger during multi-finger prehension tasks [19], [20]. Regarding the effect of grasping practice, repetitive and lifelong exposure to object manipulation with the dominant hand [21] may lead to the development of distinct force coordination patterns, which differ from those used when grasping with the non-dominant hand. The findings of Zhang et al. found that the dominant hand is more effective at controlling rapid force changes, indicating its specialization in dynamic tasks. In contrast, there was no evidence for the non-dominant hand’s specialization in stabilizing steady-state force, possibly due to the simplicity of the isometric tasks used [22]. Rearick and Santello (2002) observed that force control strategies during grasping and lifting tasks are similar across both hands, suggesting that the central nervous system employs comparable patterns for coordinating finger forces regardless of hand dominance [23]. Previous studies on finger movements, pegboard tasks, and reaching tasks suggest that the non-dominant hand may require greater control or coordination to perform tasks compared to the dominant hand. Real-life situations often involve multidigit prehension, requiring the simultaneous use of finger and thumb forces. Hence, this study aimed to compare the behavioural and neural features during lifting and grasping tasks between the dominant and non-dominant hands in healthy young adults. Based on prior understanding, it was hypothesized that the performance of the dominant hand would be better compared to the non-dominant hand. We expected this to be reflected in both neural measures (EEG band power and EEG–EMG coherence) and behavioural measures (synergy index).

## Methods

### Participants

Twenty young right-handed healthy adults with no history of neuromuscular disorders aged between 20 and 30 years (26.95 ± 2.68) participated in this study. The experimental procedure was approved by the Institute’s ethics committee at the Indian Institute of Technology, Madras (Approval number: IEC/2021-01/SKM/02/05). Written informed consent was collected from the participants in accordance with the ethical approval.

### Experimental Apparatus and Protocol

A handle was specifically designed for the experiment to measure the finger force applied by each finger. The handle was equipped with five 6 component force/torque sensors ( Nano 17, Force resolution: 0.0125 N, ATI Industrial Automation, NC, USA) as shown in Fig 1 A. Four sensors were fixed on one side of the handle to measure the fingertip forces of index, middle, ring and little fingers. The sensor to measure the thumb digit force was placed on the other side of the handle on a slider platform which can move on a rail. The total weight of the handle including the external weight of 0.25kg was 0.75kg. A displacement sensor was used to measure the position of the thumb (OADM 12U6460 sensor with resolution of 5 μm, Baumer, India). An inertial measurement unit (IMU – Model: BNO055 with a resolution of 16 bits and 2000°/s range, BOSCH, Germany), was placed on an acrylic block on top of the handle to measure the orientation of the handle. Additionally, a spirit level was incorporated on the handle, allowing the participants to visually monitor and ensure the orientation of the handle during the experiment. The experimental handle used in this study was previously employed by our research group in other experimental studies [24], [25], [26], [27].

**Fig 1.**
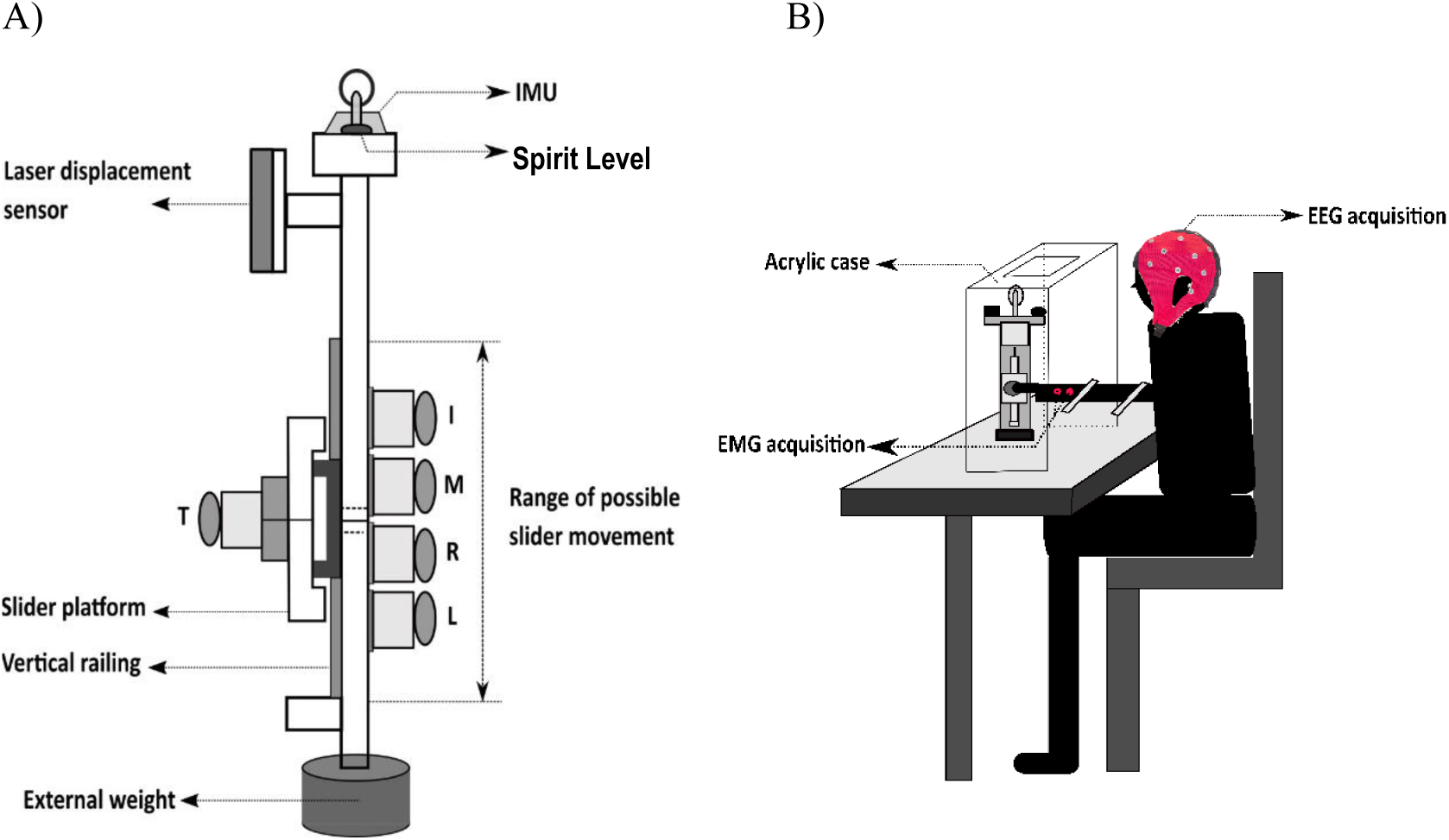
**A) Handle.** The experimental handle, made up of aluminum frame measuring 21 x 1 x 3 cm was equipped with the five force/torque sensors. An IMU sensor was mounted on an acrylic block on one side of the handle, while a spirit level was positioned on the opposite side. The positions for the five fingers were labelled as I (index), M (middle), R (ring), L (little) and T (thumb). The total weight of the handle is 0.75kg including the external weight of 0.25kg. **B) Experimental setup.** The designed handle was placed inside an acrylic case. Participants were instructed to grasp and hold the handle for initial 5 seconds gently. After 5 seconds they were asked to lift and hold the handle in the air for an additional five seconds.

A bio signal system comprising of two-channels for EMG and twenty-two channels for EEG system was developed using INTAN RHD 2216 bio amplifiers (Intan Technology, USA). The RHD 2216 amplifiers are arrays of 16 channel bipolar bio amplifiers equipped with a serial peripheral interface (SPI) on-chip and a 16-bit analog to digital converter (ADC). A sixty-four channel EEG cap (Waveguard original cap, 64-channel ANT neuro waveguard, The Netherlands; 10/10 electrode montage) was connected to the RHD amplifiers through a custom designed interface board. The EEG signals were recorded from the following 22 channels: Fp1, Fp2, AFz, Fz, F3, F4, F7, F8, Fc1, Fc2, Fc5, Fc6, Cz, C3, C4, Cp5, Cp1, Cp2, Cp6, Pz, P3, and P4. The GND electrode of the EEG cap is connected to the ground of the amplifier. The participants were instructed to clean and dry their hair before the experiment to reduce the effect of impedance. Two surface EMG signals were collected from the flexor digitorum superficialis muscle (FDS) and flexor carpi ulnaris (FCU). We have selected FDS, responsible for finger flexion and FCU, involved in wrist flexion abduction. These muscles play crucial roles in gripping and manipulating objects with multiple digits [28]. Surface electrodes in bipolar configuration were placed over the belly of the muscle with an inter electrode placement of approximately 1.5 cm [29]. A ground electrode was placed 4 to 5 cm away from the bi-polar surface electrodes [29]. The placement of electrodes was determined initially through anatomical descriptions, palpations and assessment of functional contractions [30].

### Experimental setup

The participants were instructed to sit on a chair and rest their right/left arm in a comfortable position on the table as shown in Fig 1 B. Prior to the experiment a sixty-four channel EEG cap was placed on the participants head and electrode-gel was applied to each channel using a syringe with blunt needle to reduce the electrode impedance. The electrode impedance was measured before the experiment and adjusted to be below 50kΩ.

### Experimental Protocol

The experiment consists of two task conditions: fixed and free conditions. The thumb platform was positioned on a movable base, with the task conditions defined by the mobility of the platform. In the fixed task condition, the thumb platform was securely fixed with screws between the middle and ring fingers. In the free task condition, the thumb platform was allowed to move freely; however, participants were instructed to maintain the thumb position between the middle and ring fingers. The thumb displacement was measured using a displacement measurement sensor. In the free condition, if the displacement of the thumb platform was more than 1 cm then the trial was rejected. All the participants performed the task with both the dominant hand and non-dominant hand separately. Among the twenty healthy young participants, 50% performed the task with their dominant hand and subsequently switched to the non-dominant hand, while the remaining 50% performed with their non-dominant hand before switching to the dominant hand. The trial order was randomized to control for any potential order effects. During both the fixed and free conditions, initially for a 5 second period, the participants were instructed to hold the handle gently, using all five fingers in a relaxed manner. After this initial 5 second period they were asked to lift the handle and hold it in air for another 5 seconds. The participants were instructed to maintain the handle in steady state in the lift and hold period by maintaining the bubble of the sprit level at the center. Each participant performed 25 such trials for each condition with each hand. Adequate breaks were provided between the trials and additional breaks were given upon the participants request.

### Data acquisition

The force signals from the five sensors and the displacement signal were digitized using a 16-bit NI USB 6225 and 6210 DAQs (National Instruments, Austin, TX, USA) respectively. The orientation signal from the IMU was recorded through a microcontroller. Force signals, the orientation data and the displacement data were sampled at 100 samples per second. Two-channel EMG and twenty-two channel EEG were recorded using two RHD 2216 (INTAN technologies, USA) through XEM6310 (Opal Kelly, Portland) FPGA interface board. EEG and EMG signals were sampled at 1500 Hz. EEG signal, EMG signal, finger force signal, displacement signal and the orientation data from IMU were recorded simultaneously using a custom LabView code (version 19).

### Data analysis

All the recorded signals were analysed offline in MATLAB (Version R2022a, MathWorks, USA).

### Synergy Analysis

We performed variance analysis as described by Latash et al., (2002); Zhang et al., (2009) to quantify multi finger synergy [20], [31]. An index (ΔV) of synergy was computed as the difference between the sum of the variances of the mechanical variables produced by individual digits (elemental variables) [ΣVar(EV)] and the variance of the total output of these elemental variables [Var(ΣEV)] [33]. The variance of one of the variables on the left side of equations (2), a performance variable (PV), was compared to the sum of the variances of the variables on the right side of those equations, the elemental variables (EVs), and then normalized by the latter value [19], [22], [32] (Equation 1).

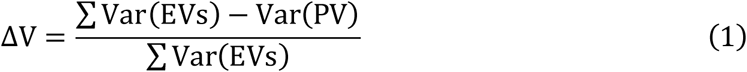

The variance analysis was conducted for performance variable, total moment of force (M_Tot_) due to the normal and tangential forces at the virtual finger (VF) and thumb (TH) level. The Virtual Finger (VF) refers to the combined action of the index, middle, ring, and little fingers, which together function as a single entity in manipulating objects. In the VF-TH level hierarchy, VF forces and moments (sum of index, middle, ring, and little fingers) and the thumb forces and moments were considered as elemental variables. At the VF-TH level, we were interested in examining the co-variation of the following quantities produced by the VF and the opposing effector: the total moment of force, M_Tot_. The total moment of force produced by the normal and tangential force should be equal to zero in equilibrium conditions.

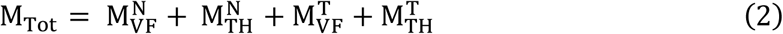

Participants were instructed to grasp and lift the handle with all five fingers positioned appropriately as demonstrated in Fig. 1. Participants were instructed to lift the handle and hold it in the air for 5 seconds, referred to as the hold period. Throughout this duration, participants were instructed to maintain the handle in a static state without tilting. To ensure this, a spirit level was employed, and the orientation of the handle was monitored using an Inertial Measurement Unit (IMU) sensor. If the tilt of the handle exceeded 3 degrees, indicating instability, the trial was rejected, and participants were asked to repeat the trial. Thus, steady-state activity was rigorously monitored and maintained.

The synergy index was computed for a two second segment of the 5-second hold period, individually for each participant, for each hand and (dominant and non-dominant hand) for each condition (Fixed and Free). Positive values of the synergy index indicate the negative covariation among the elemental variables and, a synergy stabilizing the performance variable. Fisher’s z transformation was applied to the synergy index using the equation (3) for the statistical analysis.

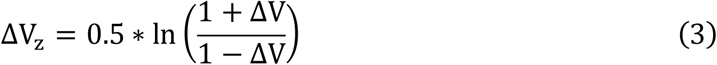

### EEG Band power

The power line interference and its harmonics from the recorded EEG signals were removed using a 50 Hz notch filter. Out of 22 channels, the following 12 EEG channels were taken for further analysis, Fc1, Fc2, Fc5, Fc6, C3, C4, Cp5, Cp1, Cp2, Cp6, P3, and P4. These 12 channels were selected based on their coverage of key brain regions involved in motor control [33], [34], [7]. During the experiment, participants were instructed to minimize eye blinks and eye movements as much as possible. This instruction aimed to reduce the occurrence of ocular motion artifacts during data collection. After recording EEG signals, we checked the signal amplitude for each trial manually. If the amplitude exceeded the threshold of 150 μV, indicating the presence of potential artifacts, the trial was rejected. We rejected 10% of the data, specifically 3 out of 28 trials, resulting in the selection of a total of 25 artifact free trials, resulting in the selection of 25 artifact-free trials. The notch filtered 12 channel EEG signals were bandpass filtered to beta band (13 – 30 Hz) using a fourth order, zero phase lag Butterworth filter. Each band signal was down-sampled to 100 Hz. The total trial duration of the task was 10 seconds.

Initially, participants were instructed to gently grasp the handle with all five fingers for 5 seconds (Relax period). After this period, they were asked to lift and hold the handle in the air for another 5 seconds (Hold period). The band power value of the EEG signal between 2 – 4 seconds of the holding phase was taken to compare the two conditions for the 12 channels mentioned above. For analysis, we focused on the middle two seconds of the 5-second hold period to minimize any minor drift that may occur during the initial and final periods.

When participants performed tasks with their right hand (dominant hand), the EEG band power of the contralateral side (left hemisphere) channels Fc1, Fc5, C3, Cp5, Cp1, and P3 were analysed. When tasks were performed with the left hand (non-dominant hand), the EEG band power of the contralateral side (right hemisphere) channels Fc2, Fc6, C4, Cp6, Cp2, and P4 were analysed. The band power of these channels were analyzed, and the contralateral corresponding channels were compared: Fc1 and Fc2, Fc5 and Fc6, C3 and C4, Cp5 and Cp6, Cp1 and Cp2, P3 and P4.

The following steps were used to calculate the Band Power - BP (21) for both the fixed and free condition.

1. The power of the samples was obtained by squaring the amplitude samples of each trial
2. Averaging the power samples across trials

The mean band power during the holding phase was averaged across all participants, and the standard error of the mean was calculated.

### EEG – EMG Coherence

The artifact removed EEG signal were bandpass filtered between 4 – 40 Hz using a fourth order, zero phase lag Butterworth filter. The notch filtered EMG signals were band pass filtered between 20 – 500 Hz using a fourth order, zero phase lag Butterworth filter. EEG and EMG signals during the lift and hold phase between 2 – 4 seconds of each trial were taken for the EEG-EMG coherence calculation. EMG signals were not rectified prior to calculating corticomuscular coherence (CMC). Coherence gives a linear relationship between two signals in the frequency domain [35]. Coherence is obtained through the normalization of the cross-spectrum of the signals x(t) and y(t) [36].

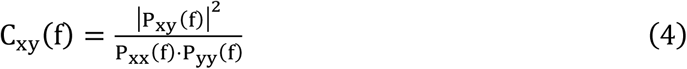

Where, P_xx_ (f) is the power spectrum of the time series signal x(t), and P_yy_ (f) is the power spectrum of the time series signal y(t).

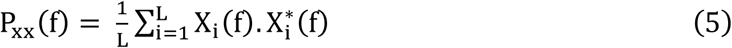

where X_i_ (f) is the Fourier transform of the i-th signal segment within the total sample of L, where * denotes the complex conjugate. P_xy (f) is the cross spectrum of the two signals x (t) and y(t) is indicated as,

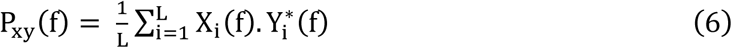

where Y_i_ (f) is the Fourier transform of the signal segment I of the total sample of L and * represents the complex conjugate. Corticomuscular coherence characterizes the coherence of the cortical signals x(t) and the muscle signals y(t).

Coherence was calculated between each EEG channel (14 channels) and each EMG channel (FDS and FCU) using the mscohere function in MATLAB’s Signal Processing Toolbox. For each trial, the data segment of 3000 samples was divided into 8 non-overlapping segments, with a Hamming window applied to each segment. This segmentation provided a window length of 375 samples (equivalent to 250 ms) and a frequency resolution of 4 Hz. The choice of non-overlapping segments was made to ensure independent estimates across segments. Coherence estimates were averaged across these segments to increase the reliability of the measure. The raw coherence values were then transformed into z-scores using the formula [37], [38] (4),

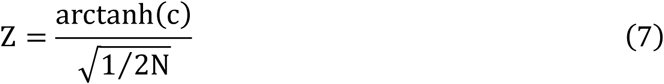

where, N=8 denote the number of segments. The 95% confidence intervals (CI) for coherence values were calculated based on non-overlapping segments, using the formula [37], [38], [39](5)

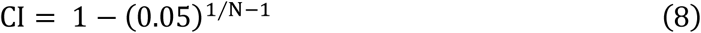

The 95% confidence interval (CI) for coherence values, with N=8, is approximately 0.348. This means that coherence values above this threshold can be considered significantly different from zero at the 95% confidence level. Mean squared coherence was calculated between 2 – 4 seconds of the holding phase of the task for the two conditions. For analysis, we focused on the middle two seconds of the 5-second hold period to minimize any minor drift that may occur during the initial and final periods. The mean squared coherence was averaged across the trials, across the participants and standard error of mean was calculated.

## Statistical analysis

All statistical analysis were performed in R. A two-way repeated measures ANOVA was conducted with factors: Conditions (2 conditions: Fixed and Free) X Hand (2 hands: DH and NDH) on EEG band power, synergy index and EEG–EMG coherence. This analysis aimed to examine the changes in EEG band power, synergy index and EEG – EMG coherence. The significant level p < 0.05 was chosen for the statistical analysis. Partial eta-squared (η^2^) was stated as effect size. The sphericity test was executed on the data for all the cases and Huynh-Feldt (H-F) criterion was applied as necessary to adjust the degree of freedom. Two one sided t test (TOST) analysis was performed on the EEG band power, synergy index and EEG – EMG coherence between the two task conditions with the smallest effect size of interest (SESOI = 1.04) as equivalence bounds. Equivalence bound of ΔV = 1.04 was chosen as lower and upper bound for the statistical power of 95% [40].

## Results

### Synergy index of total moment of force

There was no difference observed in the synergy index between the left hand and right (DH and NDH of the right handers) under both task conditions. Since there was no difference observed, TOST analysis was performed by considering the upper and lower bound ( ΔU and ΔL) as ±1.04 between the C3 and C4 channel EEG band power. TOST analysis showed a statistical equivalence on the EEG band power between the C3 and C4 channel as shown in Fig 2. However, synergy index of the fixed condition was higher than the free condition in both dominant and non-dominant hands. This was supported by a two-way repeated ANOVA measures by considering the factors conditions and hands. Two-way repeated ANOVA measures revealed a main effect only for the factor conditions (F_(1,19)_ = 81.08, η² = 0.81, p < 0.0001). However, no main effect was observed for the factor hands. The hands x condition interaction was not statistically significant.

**Fig 2.**
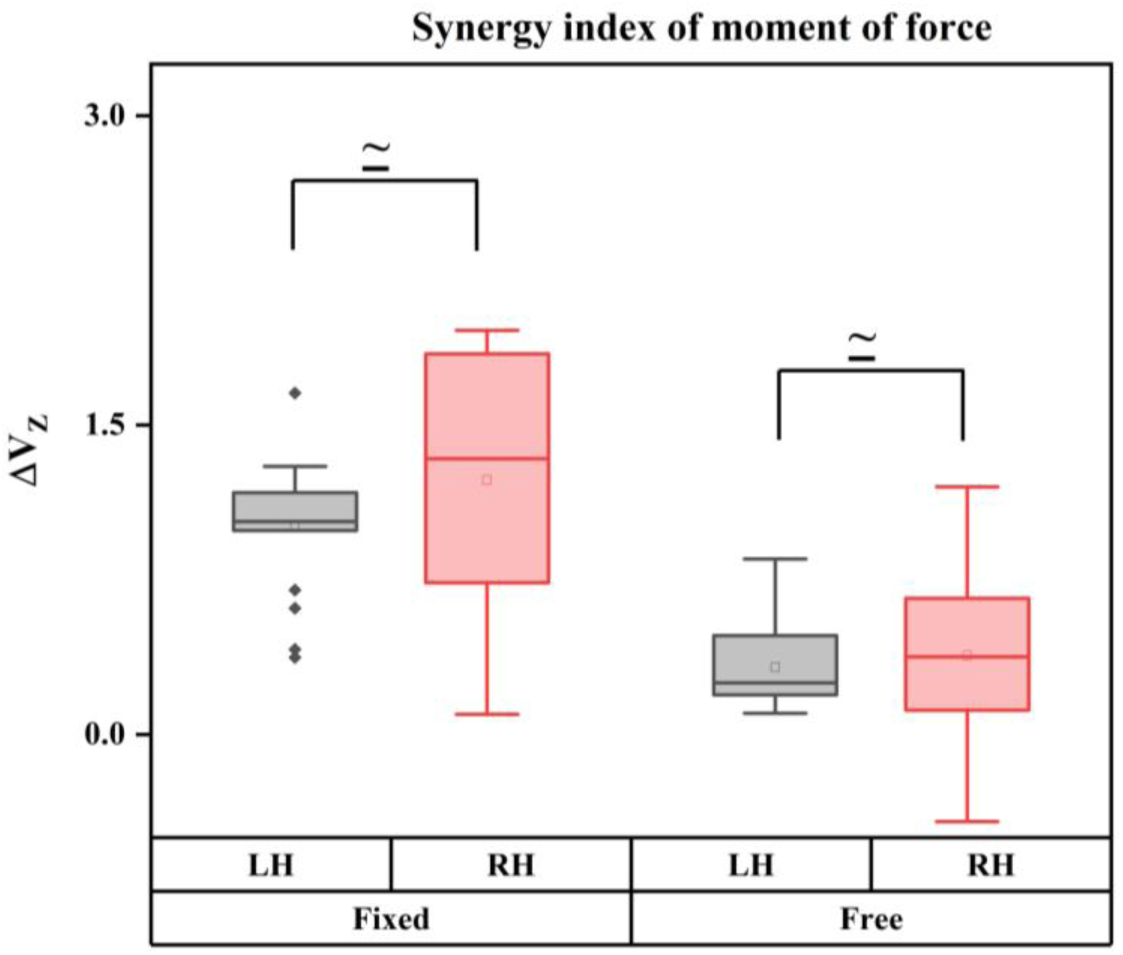
Synergy index of moment of force. There was no difference observed in synergy index between the hands in both task conditions. TOST analysis showed statistical equivalence of the synergy index of LH and RH (DH and NDH of the right handers). However, synergy index of moment of force in the fixed condition was higher than the free condition for both hands (p < 0.0001).

### EEG band power

There was no difference observed between the C4 and C3 EEG band power corresponding to the contralateral channel in both task conditions. Since there was no difference observed, TOST analysis was performed by considering the upper and lower bound ( ΔU and ΔL) as ±1.04 between the C3 and C4 channel EEG band power. TOST analysis showed a statistical equivalence on the EEG band power between the C3 and C4 channel as shown in Fig 3. However, EEG band power of the fixed condition was higher than the free condition in both dominant and non-dominant hands. This was supported by a two-way repeated ANOVA measures by considering the factors conditions and hands. ANOVA results revealed a main effect only for the factor task conditions (F_(1,19)_ = 11.07, η² = 0.37, p < 0.03). Nevertheless, no main effect was not observed for the factor hands. The hands x conditions interaction was also not statistically significant.

**Fig 3.**
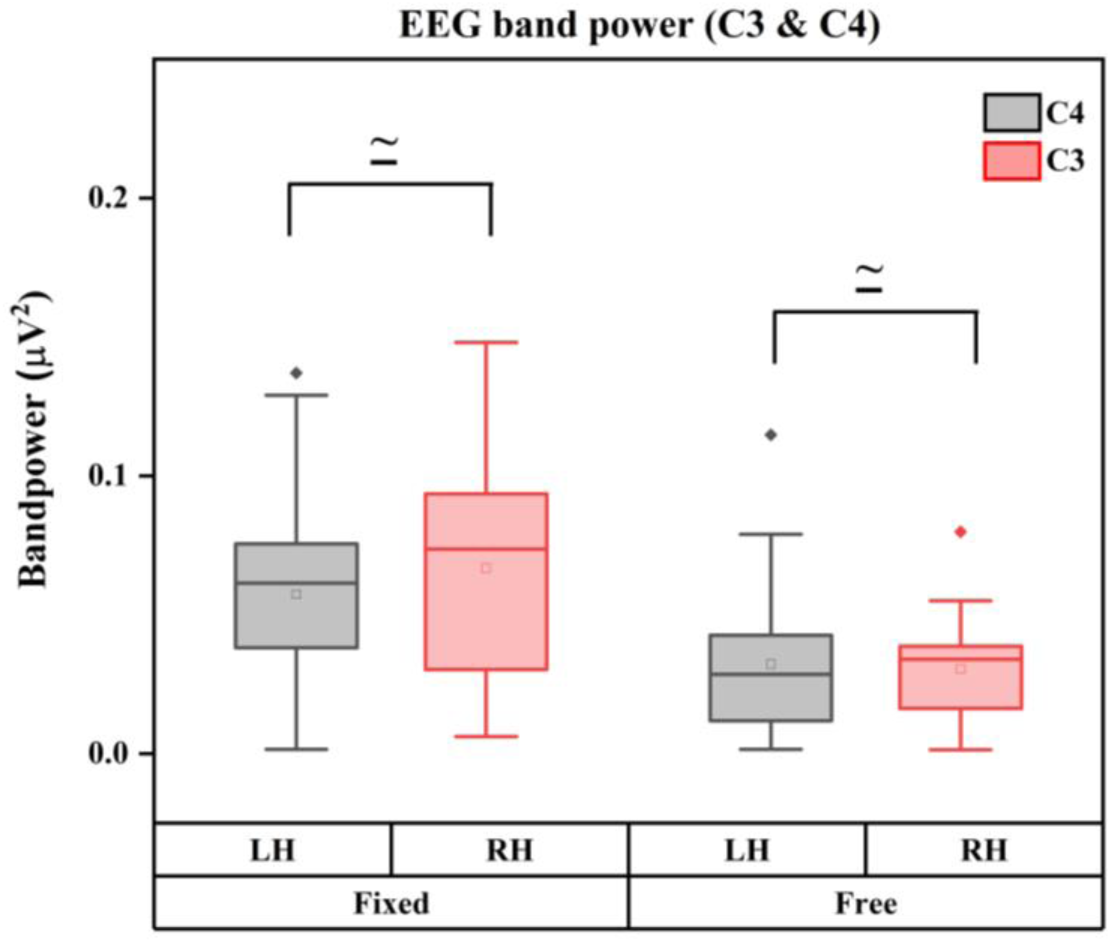
EEG band power. There was no difference observed in EEG band power between the channel C3 and C4 in both task conditions. TOST analysis showed statistical equivalence of the EEG band power of C4 and C3 activities corresponding to the contralateral hand in both fixed and free task conditions. However, EEG band power of fixed condition was higher than the free condition for both hands (p < 0.05).

### EEG – EMG coherence

The magnitude of EEG–EMG coherence between the hands exhibited comparable values under both task conditions. Therefore, TOST analysis was performed by considering the upper and lower bound (ΔU and ΔL) as ±1.04 between the left and right hand for both task conditions. TOST analysis showed a statistical equivalence on the EEG – EMG coherence between the hands in both task conditions as shown in Fig 4. However, magnitude of EEG – EMG coherence of the fixed condition was higher than the free condition in both left and right hands. This was supported by a two-way repeated ANOVA measures by considering the factors conditions and hands of the EEG – EMG coherence. ANOVA results revealed a main effect only for the factor task conditions (F_(1,19)_ = 26.17, η² = 0.49, p < 0.0001). However, no main effect was observed for the factor hands. The hands x conditions interaction was also not statistically significant.

**Fig 4.**
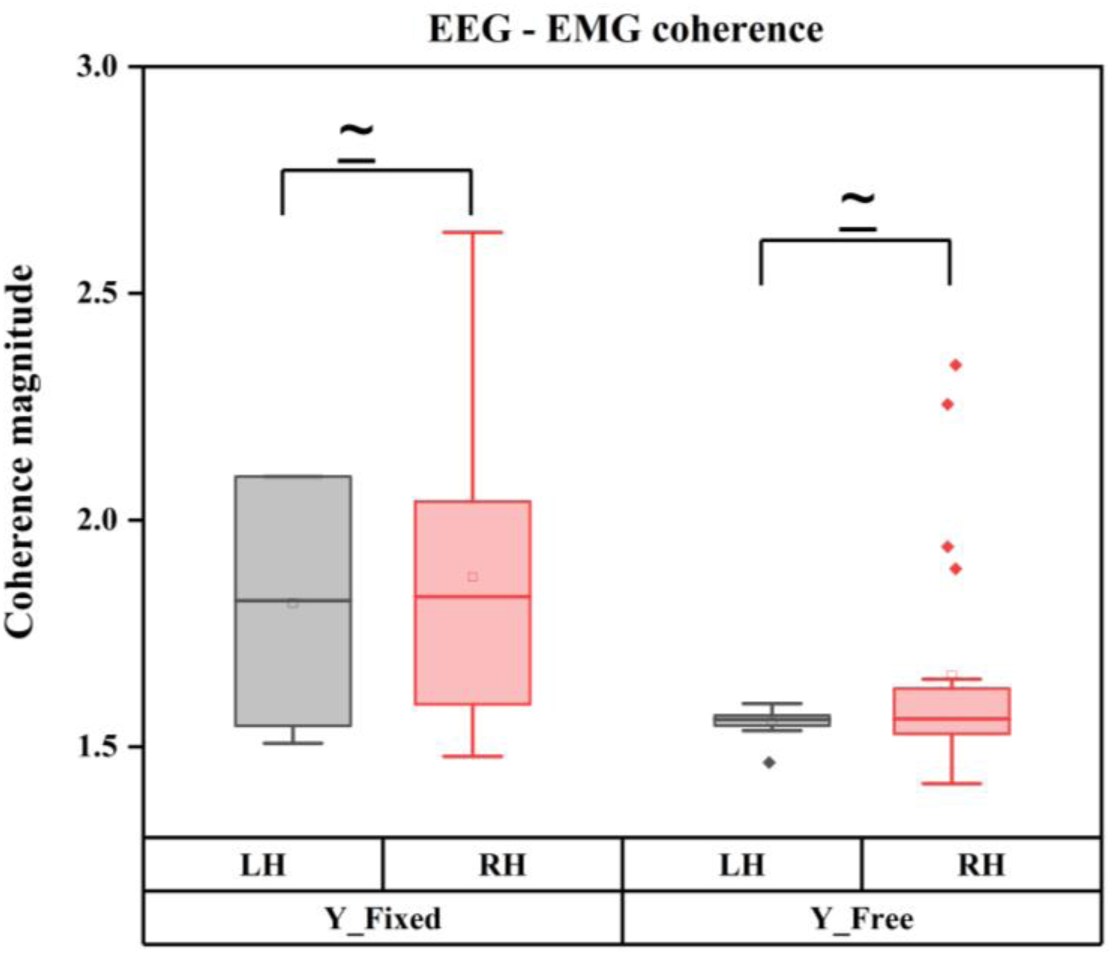
EEG – EMG coherence. The magnitude of EEG – EMG coherence was comparable between the hands in both task conditions. TOST analysis revealed a statistical equivalence between the left and right hand in both task conditions. However, EEG – EMG coherence was higher in the fixed condition compared to the free condition (p < 0.0001).

## Discussion

The primary aim of this study was to compare the behavioural and neural features during lifting and grasping tasks between the dominant and non-dominant hands in healthy young adults. It was hypothesized that the performance of the dominant hand would be better compared to the non-dominant hand. We expected this to be reflected in both neural measures (EEG band power and EEG–EMG coherence) and behavioural measures (synergy index), with more pronounced results for the dominant hand. However, our results contradicted the hypothesis, as no differences were observed in synergy index, EEG band power, and EEG–EMG coherence, between the dominant and non-dominant hands. These findings suggest that the non-dominant hand can function similarly to the dominant hand for the given task. The implications of these findings are discussed in the following paragraphs.

In our study, participants had to maintain the handle in a steady state during both task conditions. The fixed task condition is similar to isometric tasks, where constant force is required to keep the handle steady. However, in the free task condition, the thumb platform is freely movable due to the frictionless setup, which introduces a level of dynamic interaction. The results indicate a clear task-level difference in the synergy index for both dominant and non-dominant hand, suggesting that the modulation of force coordination varies between the two task conditions. Despite the hypothesized differences, the similar synergy indices observed between the dominant and non-dominant hands suggest that both hands are capable of achieving comparable levels of coordination and stability during the task. This lack of difference in the synergy index aligns with studies on handling fragile objects with both hands [41] and isometric conditions [22]. In isometric conditions, muscle contraction occurs without changing muscle (muscle + tendon) length. Performance differences typically associated with hand used in dynamic reaching tasks may not be evident in this task, as it does not involve complex movements or dynamic adjustments.

These results suggest that, despite the differing nature of the tasks, the underlying motor control mechanisms may be similarly effective across both hands. Notably, the non-dominant hand demonstrated a level of performance comparable to the dominant hand, even under the dynamic thumb condition which was supported by TOST analysis. This finding is particularly intriguing, as the dynamic condition introduces greater complexity in thumb control due to the need for constant adjustments. The ability of the non-dominant hand to match the performance of the dominant hand in this condition suggests that the cortical activity and finger force coordination are not exclusively reliant on hand used. Instead, both hands appear to utilize similar adaptive strategies to achieve coordinated control of finger forces across varying task demands. These results highlight the robustness of the motor control system in facilitating task-specific adjustments in finger force coordination, regardless of hand being used. Furthermore, this suggests that the non-dominant hand can compensate effectively, potentially reducing reliance on the dominant hand, even in tasks requiring fine motor adjustments.

A possible explanation for this lack of observed differences lies in the different roles of the visual and motor systems during the task. Reaching tasks require significant coordination between these systems, but after grasping and lifting the handle in both task conditions, their involvement reduces. This reduced engagement of the visual system might elucidate the lack of differences observed between the hands in both task conditions in our study. These results are partially consistent with another study from our group, where finger force synergies were similar between hands in non-challenging tasks, while the dominant hand showed better coordination during more challenging tasks. However, the task in the above-mentioned study also involved continuous monitoring of the thumb platform displacement, which might have influenced the observed outcomes [27].

However, our findings contradict the dynamic dominance hypothesis, which suggests that the dominant arm uses more sophisticated neural control mechanisms and coordination strategies. According to this hypothesis, differences in muscle torques and hand path trajectories should be evident between the dominant and non-dominant hands [1], [3], [4]. The absence of such differences in the present study suggests that the task demands may not have been sufficiently complex or dynamic to elicit dominance-related disparities. Alternatively, these findings might indicate that, for tasks involving steady-state force maintenance and limited visual-motor engagement, both hands are capable of achieving comparable performance and stability. This interpretation is further supported by the EEG results, which revealed comparable cortical activity in the contralateral channels for both hands in both task conditions. Additionally, no differences were observed in EEG band power between the hands in either task condition. These findings imply that the cortical control mechanisms involved in the task are similar for both hands, regardless of the hand used. The comparable EEG band power suggests that both hands employ equivalent neural resources to meet the demands of the task, further underscoring the uniformity in their performance. The lack of observed differences between the hands in EEG band power may also reflect the task’s emphasis on steady-state force maintenance rather than dynamic adjustments or fine motor precision. Tasks involving greater variability or rapid transitions might engage the cortex differently and reveal dominance-related differences. The findings of this study contradict the expected differences based on the dominant hemisphere’s known anatomical and functional advantages [10]. The hemispheric differences suggested by previous findings could be related to the specialized neural mechanisms for skilled movements in the dominant hand. However, EEG band power in the contralateral channels corresponding to hand movement (C3 for the right hand and C4 for the left hand) was considerably higher in the fixed condition compared to the free condition. This observation aligns with the findings on finger force coordination and suggests task-level modulation in cortical activity specific to the contralateral hand.

Moreover, the similar corticomuscular coherence (CMC) magnitude observed between the dominant and non-dominant hands further supports the notion of comparable performance of the hands. Despite differences in hemispheric representation for the dominant and non-dominant hands, the overall performance and functionality of the hands did not differ considerably. These findings align with previous studies, which observed no differences between the two hands during the steady-state phase [17]. The similarity in EEG band power across both hands suggests that equivalent neural resources are engaged, regardless of hand dominance. This uniform cortical activity likely supports the observed comparable EEG–EMG coherence, as the synchronization of cortical and muscle activity depends on the consistency and efficacy of cortical signals reaching the muscles.

Additionally, task-level modulations in corticomuscular coherence (CMC) were observed for both hands. In the free condition, the altered friction between the platform and the thumb is likely perceived by proprioceptors and cutaneous receptors in the fingertip. This sensory input prompts adjustments in grip and tangential forces, allowing the fingers to maintain the handle’s equilibrium. These adjustments demonstrate how finger coordination adapts dynamically to meet task-specific requirements [26]. Previous studies demonstrated that synergistically activated muscles are driven by a shared beta-band cortical drive, which can be quantified using intramuscular coherence [14]. The observed reduction in CMC during the free condition can thus be interpreted as a decrease in finger coordination due to changes in friction. This reduction likely reflects a shift in neural control strategies, where the need for finer adjustments to changing friction demands results in less synchronous cortical input to the muscles. These results align with previous findings, where beta-band CMC was reduced during tasks involving object compliance. Specifically, studies demonstrated that CMC was lower when using a compliant object like a spring compared to a rigid object such as a wooden dowel during a two-finger pinch grasp [37]. Additionally, differences in the neural and behavioural features between the task conditions in the dominant hand of young adults have been extensively discussed in our previous findings [26].

Previous studies have suggested that interhemispheric monitoring and performance evaluation play a key role in motor control and hand dominance, with hemispheric interactions influencing how each hand’s performance is monitored and adjusted [42], [43]. However, the coordination and force control required in our tasks may necessitate similar motor strategies from both hands, enabling them to perform the task effectively regardless of dominance. Taken together, task-level modulation was observed between the task conditions in both hands, specifically in cortical activity, finger force coordination, and EEG-EMG coherence. The lack of differences in synergy indices, combined with the comparable neural features observed across task conditions, underscores the adaptability of motor control strategies and the uniformity in how both hands respond to task demands. The lack of interaction effect between the hand and conditions signify that the hands adjust to task demands in a non-different manner. These findings highlight the capacity of both hands to achieve comparable performance in tasks requiring steady-state force control, despite their typical dominance-related roles in other contexts.

## Conclusion

In this study we compared the behavioural and neural features between the dominant hand and non-dominant hand during grasping and lifting tasks under two task conditions. Specifically, we compared the synergy index and EEG band power, EEG-EMG coherence as behavioural and neural features. Our study revealed that there was no difference observed between the dominant and non-dominant hands in both behavioural and neural features. In conclusion, these findings indicate that both the dominant and non-dominant hands can achieve comparable performance levels in the given tasks, despite the dominant hand’s typical advantages in dynamic motor tasks. Task-dependent modulations were observed, particularly in the free condition, where variations in friction led to adjustments in both behavioural and neural features. Further studies should explore how different task complexities and dynamic conditions might reveal more pronounced differences between the hands.

## Acknowledgement

This work was supported by the American Express Lab for Data Analytics, Risk and Technology, IIT Madras, vide reference no SB20210346CPAMEXAMEHOC (awarded to Varadhan SKM). The funders played no role in the experimental design, protocol, data collection, analysis and in the manuscript preparation. We thank Thomas Jacob for his valuable contribution to the data collection process.

## Data availability

The authors do not report data and therefore the pre-registration and data availability requirements are not applicable.

## Authors Contribution

Data collection – B.E; Methodology – B.E., V.S.K.M., S.B.; Formal analysis – B.E.; Writing original draft – B.E.; Writing review, Review and Editing – B.E., V.S.K.M., S.B.

